# Low coverage whole genome sequencing enables accurate assessment of common variants and calculation of genome-wide polygenic scores

**DOI:** 10.1101/716977

**Authors:** Julian R. Homburger, Cynthia L. Neben, Gilad Mishne, Alicia Y. Zhou, Sekar Kathiresan, Amit V. Khera

## Abstract

**Background:** The inherited susceptibility of common, complex diseases may be caused by rare, ‘monogenic’ pathogenic variants or by the cumulative effect of numerous common, ‘polygenic’ variants. As such, comprehensive genome interpretation could involve two distinct genetic testing technologies -- high coverage next generation sequencing for known genes to detect pathogenic variants and a genome-wide genotyping array followed by imputation to calculate genome-wide polygenic scores (GPSs). Here we assessed the feasibility and accuracy of using low coverage whole genome sequencing (lcWGS) as an alternative to genotyping arrays to calculate GPSs.

**Methods:** First, we performed downsampling and imputation of WGS data from ten individuals to assess concordance with known genotypes. Second, we assessed the correlation between GPSs for three common diseases -- coronary artery disease (CAD), breast cancer (BC), and atrial fibrillation (AF) -- calculated using lcWGS and genotyping array in 184 samples. Third, we assessed concordance of lcWGS-based genotype calls and GPS calculation in 120 individuals with known genotypes, selected to reflect diverse ancestral backgrounds. Fourth, we assessed the relationship between GPSs calculated using lcWGS and disease phenotypes in 11,502 European individuals seeking genetic testing.

**Results:** We found imputation accuracy r^2^ values of greater than 0.90 for all ten samples -- including those of African and Ashkenazi Jewish ancestry -- with lcWGS data at 0.5X. GPSs calculated using both lcWGS and genotyping array followed by imputation in 184 individuals were highly correlated for each of the three common diseases (r^2^ = 0.93 - 0.97) with similar score distributions. Using lcWGS data from 120 individuals of diverse ancestral backgrounds, including South Asian, East Asian, and Hispanic individuals, we found similar results with respect to imputation accuracy and GPS correlations. Finally, we calculated GPSs for CAD, BC, and AF using lcWGS in 11,502 European individuals, confirming odds ratios per standard deviation increment in GPSs ranging 1.28 to 1.59, consistent with previous studies.

**Conclusions:** Here we show that lcWGS is an alternative approach to genotyping arrays for common genetic variant assessment and GPS calculation. lcWGS provides comparable imputation accuracy while also overcoming the ascertainment bias inherent to variant selection in genotyping array design.

## BACKGROUND

Cardiovascular disease and cancer are common, complex diseases that remain leading causes of global mortality [1]. Long recognized to be heritable, recent advances in human genetics have led to consideration of DNA-based risk stratification to guide prevention or screening strategies. In some cases, such conditions can be caused by rare, ‘monogenic’ pathogenic variants that lead to a several-fold increased risk -- important examples are pathogenic variants in *LDLR* that cause familial hypercholesterolemia and pathogenic variants in *BRCA1* and *BRCA2* that underlie hereditary breast and ovarian cancer syndrome. However, the majority of individuals afflicted with these diseases do not harbor any such pathogenic variants. Rather, the inherited susceptibility of many complex traits and diseases is often ‘polygenic,’ driven by the cumulative effect of numerous common variants scattered across the genome [2].

Genome-wide polygenic scores (GPSs) provide a way to integrate information from numerous sites of common variation into a single metric of inherited susceptibility and are now able to identify individuals with a several-fold increased risk of common, complex diseases, including coronary artery disease (CAD), breast cancer (BC), and atrial fibrillation (AF) [3]. For example, for CAD, we noted that 8% of the population inherits more than triple the normal risk on the basis of polygenic variation, a prevalence more than 20-fold higher than monogenic familial hypercholesterolemia variants in *LDLR* that confer similar risk [3].

Comprehensive genome interpretation for common, complex disease therefore could involve both high-fidelity sequencing of important driver genes to identify potential monogenic risk pathogenic variants and a survey of all common variants across the genome to enable GPS calculation. High coverage whole genome sequencing (hcWGS; for example, 30X coverage) will likely emerge as a single genetic testing strategy, but current prices remain a barrier to large-scale adoption. Instead, the traditional approach has mandated use of two distinct genetic testing technologies -- high coverage next generation sequencing (NGS) of important genes to detect pathogenic variants and a genome-wide genotyping array followed by imputation to calculate GPSs.

Low coverage whole genome sequencing (lcWGS; for example, 0.5X coverage) followed by imputation is a potential alternative approach to genotyping arrays for assessing the common genetic variants needed for GPS calculations. Several recent studies have demonstrated the efficiency and accuracy of lcWGS for other applications of statistical genetics, including local ancestry deconvolution, complex trait association studies, and detection of rare genetic variants [4–7].

We developed a pipeline for common genetic variant imputation using lcWGS data on samples from the 1000 Genomes Project (1KGP) and Genome in a Bottle (GIAB) Consortium and herein demonstrate imputation accuracy for lcWGS similar to genotyping arrays. Using three recently published GPSs for CAD [3], BC [8], and AF [3], we show high technical concordance in GPSs calculated from lcWGS and genotyping arrays. Finally, using our pipeline in a large European population seeking genetic testing, we observe similar GPS risk stratification performance as previously published array-based results [3,8].

## METHODS

### Study design

The study design is summarized in Figure 1 and described in detail below. The pipeline validation data set (n = 10) was used to assess imputation accuracy for common genetic variants (Figure 1A). The technical concordance cohort (n = 184) was used to assess the correlation between three previously published GPSs for CAD [3], BC [8], and AF [3] from lcWGS and genotyping arrays (Figure 1B). The diverse ancestry data set (n = 120) was used to assess imputation accuracy for common genetic variants and performance of GPS_CAD_, GPS_BC_, and GPS_AF_ (Figure 1B). The clinical cohort (n = 11,502) was used to assess performance of GPS_CAD_, GPS_BC_, and GPS_AF_ in a large European population seeking genetic testing (Figure 1B).

**Figure 1.**
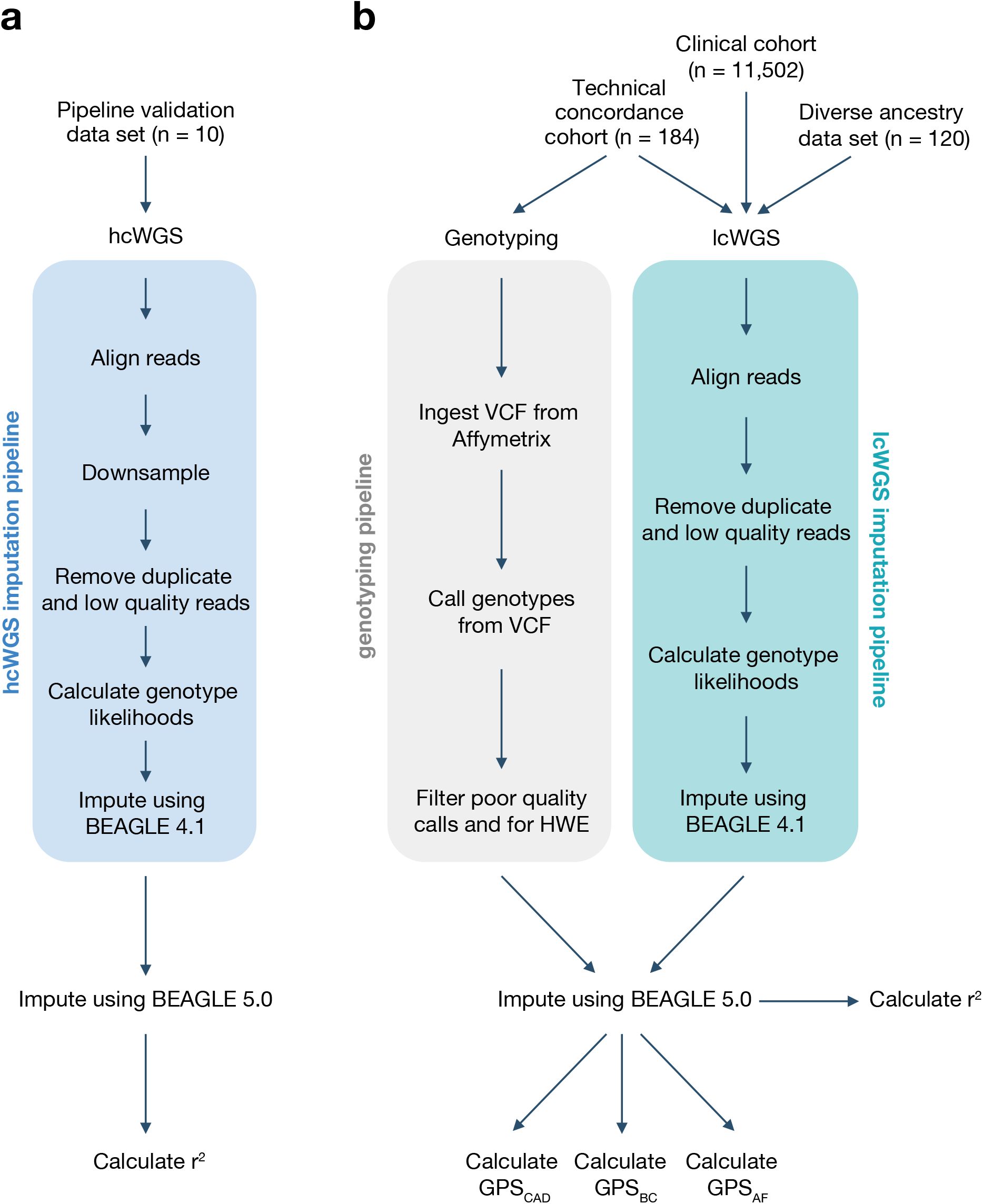
Study design and imputation pipelines. The study design has four groups: (A) pipeline validation data set and (B) technical concordance cohort, diverse ancestry data set, and clinical cohort. The imputation pipeline for each group is depicted. hcWGS, high coverage whole genome sequencing. lcWGS, low coverage whole genome sequencing. HWE, Hardy–Weinberg equilibrium. GPS, genome-wide polygenic score. CAD, coronary artery disease. BC, breast cancer. AF, atrial fibrillation.

### Data set and cohort selection

The pipeline validation data set included seven globally representative samples from 1KGP populations (HG02155, NA12878, HG00663, HG01485, NA21144, NA20510, and NA19420; see Supplementary Table 1, Additional File 1) and a trio of Ashkenazi samples (NA24385, NA24143, and NA24149) from the GIAB Consortium (Figure 1A).

The technical concordance cohort included DNA samples from 184 individuals whose healthcare provider had ordered a Color multi-gene panel test (Figure 1B). All individuals 1) had 85% or greater European genetic ancestry calculated using fastNGSadmix [9] using 1KPG as the reference panel, 2) self-identified as ‘Caucasian’, and 3) did not have pathogenic or likely pathogenic variants in the multi-gene NGS panel test, as previously described [10] (see Supplementary Methods, Additional File 2). Demographics are provided in Supplementary Table 2, Additional File 1. All phenotypic information was self-reported by the individual through an online, interactive health history tool. Of the 184 individuals, 61 individuals reported having a personal history of CAD (defined here as a myocardial infarction or coronary artery bypass surgery), 62 individuals reported no personal history of CAD, and 61 individuals reported no personal history of CAD but were suspected to have a high GPS_CAD_ based on preliminary analysis. This preliminary analysis included imputation from multi-gene panel and off-target sequencing data, which has been shown to have similar association statistics and effect sizes compared to genotyping arrays [4]. These individuals were included in the technical concordance cohort to artificially create a relatively uniform distribution of GPS_CAD_ in the data set. Correlation coefficients between GPS_CAD_ from lcWGS and genotyping array were calculated after removing the 61 individuals who were suspected to have a high GPS_CAD_ based on multi-gene panel and off-target sequencing data to avoid artificial inflation of the correlation coefficient. Two individuals who reported no personal history of CAD but were suspected to have a high GPS_CAD_ failed genotyping (quality control call rate of < 97%) and lcWGS (overall coverage of < 0.5X), leaving a total of 182 individuals for analyses.

The diverse ancestry data set included a total of 120 samples from the following populations from 1KGP: Han Chinese in Beijing, China (CHB); Yoruba in Ibadan, Nigeria (YRI); Gujarati Indian from Houston, Texas (GIH); Americans of African Ancestry in Southwest USA (ASW); Mexican Ancestry from Los Angeles, USA (MXL); and Puerto Ricans from Puerto Rico (PUR) (see Supplementary Table 3, Additional File 1; Figure 1B). Four samples, including NA18917 and NA19147 from the YRI population and NA19729 and NA19785 from the MXL population, were below the target 0.5X coverage and removed from analyses.

The clinical cohort included DNA samples from 11,502 individuals whose healthcare provider had ordered a Color multi-gene panel test (Figure 1B). All individuals 1) had 90% or greater European genetic ancestry calculated using fastNGSadmix [9] using 1KPG as the reference panel, 2) self-identified as ‘Caucasian’, 3) provided history of whether they had a clinical diagnosis of CAD, BC, or AF, and 4) did not have pathogenic or likely pathogenic variants detected in the multi-gene NGS panel test, as previously described [10] (see Supplementary Methods, Additional File 2). Demographics are provided in Supplementary Table 2, Additional File 1. All phenotypic information was self-reported by the individual through an online, interactive health history tool.

### Whole genome sequencing

DNA was extracted from blood or saliva samples and purified using the Perkin Elmer Chemagic DNA Extraction Kit (Perkin Elmer, Waltham, MA) automated on the Hamilton STAR (Hamilton, Reno, NV) and the Chemagic Liquid Handler (Perkin Elmer, Waltham, MA). The quality and quantity of the extracted DNA were assessed by UV spectroscopy (BioTek, Winooski, VT). High molecular weight genomic DNA was enzymatically fragmented and prepared using the Kapa HyperPlus Library Preparation Kit (Roche Sequencing, Pleasanton, CA) automated on the Hamilton Star liquid handler and uniquely tagged with 10 bp dual-unique barcodes (IDT, Coralville, IA). Libraries were pooled together and loaded onto the NovaSeq 6000 (Illumina, San Diego, CA) for 2 × 150 bp sequencing.

For the pipeline validation data set, all samples underwent WGS with mean coverage of 13.22X (range 7.82X to 17.30X); downsampling was then performed using SAMtools to simulate lcWGS. For the technical concordance cohort, all samples underwent lcWGS with mean coverage of 1.24X (range 0.54X to 1.76X). Imputed genotypes were compared with published, high-confidence known genotypes from 1KGP and the GIAB Consortium. For the diverse ancestry data set, all samples underwent lcWGS with mean coverage of 0.89X (range 0.68X to 1.24X). For the clinical cohort, all samples underwent lcWGS with mean coverage of 0.95X (range 0.51X to 2.57X).

### Downsampling

For the pipeline validation data set, aligned reads were downsampled using SAMtools [11] to 2.0X, 1.0X, 0.75X, 0.5X, 0.4X, 0.25X, and 0.1X coverage. For the technical concordance cohort, aligned reads were downsampled to 1.0X, 0.75X, 0.5X, 0.4X, 0.25X, and 0.1X coverage. In a few cases in the technical concordance cohort, the primary samples had fewer reads than the target downsample. In those situations, all of the reads were retained. For example, if the primary sample only had 0.8X coverage, when downsampled to 1.0X, all reads were retained. Downsampling was repeated using two independent seeds in SAMtools. Once the downsampled data was generated, the imputation was repeated to generate imputed genotypes using only the downsampled reads.

### Imputation site selection

All data sets and cohorts were imputed to a set of autosomal SNP and insertion-deletion (indel) sites from 1KGP with greater than 1% allele frequency in any of the five 1KGP super populations (African, American, East Asian, European, and South Asian), for a total of 21,770,397 sites. This is hereafter referred to as the ‘imputation SNP loci.’ Multi-allelic SNPs and indels were represented as two biallelic markers for imputation.

### Genotype likelihood calculations and imputation

Genotype likelihood calculations and imputation were performed independently for each sample. Sequence reads were aligned with the human genome reference GRCh37.p12 using the Burrows-Wheeler Aligner (BWA) [12], and duplicate and low quality reads were removed. Genotype likelihoods were then calculated at each of the biallelic SNP loci in the imputation SNP loci that were covered by one or more sequencing reads called using the mpileup command implemented in bcftools version 1.8 [13]. Indels or multi-allelic sites were not included in this first genotype likelihood calculation. Reads with a minimum mapping alignment quality of 10 or greater and bases with a minimum base quality of 10 or greater were included. Genotype likelihoods at each observed site were then calculated using the bcftools call command with allele information corresponding to the imputation SNP loci. This procedure discarded calls with indels or calls where the observed base did not match either the reference or expected alternate allele for the SNP locus.

Imputation was performed using the genotype likelihood imputation option implemented in BEAGLE 4.1 [14]. This imputation used default parameters except with a model scale parameter of 2 and the number of phasing iterations to 0. A custom reference panel was constructed for each sample being imputed by selecting the 250 most similar samples to that sample from 1KGP Phase 3 release using Identity-by-State (IBS) comparison. A reference panel size of 250 was selected to best balance imputation run time and accuracy (see Supplementary Figure 1, Additional File 2). To ensure that IBS values were comparable across samples, a set of regions consistently sequenced at high depth (> 20X) across all samples was utilized. When imputation was performed on samples included in 1KGP Phase 3 release, that sample and any related samples were excluded from the custom reference panel.

To generate genotypes at all of the remaining untyped sites, a second round of imputation was performed using BEAGLE 5.0 [15]. This imputation used default settings and included the full 1KGP as the imputation reference panel. To note, when performing analysis using 1KGP samples, any related individuals were removed. Each sample then had imputed genotype calls at each of the imputation SNP loci. Indels and multiallelic sites were included in this second genotype likelihood calculation.

### Genotyping array

DNA was extracted from blood or saliva samples and purified using the Perkin Elmer Chemagic DNA Extraction Kit (Perkin Elmer, Waltham, MA) automated on the Hamilton STAR (Hamilton, Reno, NV) and the Chemagic Liquid Handler (Perkin Elmer, Waltham, MA). The quality and quantity of the extracted DNA were assessed by UV spectroscopy (BioTek, Winooski, VT).

DNA was genotyped on the Axiom UK Biobank Array by Affymetrix (Santa Clara, CA). Genotypes were filtered according to the manufacturer’s recommendations, removing loci with greater than 5% global missingness and those that significantly deviated from Hardy-Weinberg equilibrium. In addition, all A/T and G/C SNPs were removed due to potential strand inconsistencies. Each of the remaining SNPs were aligned with the hg19 reference sequence to correctly code the reference alleles as allele 1, matching the sequencing data.

To generate genotypes at all of the remaining untyped sites, imputation was performed using BEAGLE 5.0 [15]. This imputation used default settings and included the full 1KGP as the imputation reference panel. To note, when performing analysis using 1KGP samples, any related individuals were removed. Each sample then had imputed genotype calls at each of the imputation SNP loci.

### Imputation accuracy and quality assessment

Imputation accuracy for 1KGP and GIAB samples was calculated by comparing imputation results with previously released genotypes, excluding regions marked as low confidence by GIAB.

Imputation accuracy on the genotyped samples was assessed on 470,363 sites that were included on the genotyping array at different allele frequency buckets: 257,362 sites with greater than 5% allele frequency, 119,978 sites between 1-5% allele frequency, and 93,022 sites with less than 1% allele frequency. Imputation quality was assessed through site-specific dosage r^2^ comparing with genotype values from the genotyping array.

### GPS selection

The GPSs for CAD [3], BC [8], and AF [3] were previously published and selected based on their demonstrated ability to accurately predict and stratify disease risk as well as identify individuals at risk comparable to monogenic disease. GPS_CAD_ contained 6,630,150 polymorphisms, GPS_BC_ contained 3,820 polymorphisms, and GPS_AF_ contained 6,730,541 polymorphisms. All loci included in these scores were included in the imputation SNP loci.

### GPS normalization

In the clinical cohort, raw GPSs were normalized by taking the standardized residual of the predicted score after correction for the first 10 principal components (PC) of ancestry [16]. PCs were calculated by projecting lcWGS samples into 10 dimensional PC analysis (PCA) space using the LASER program [17]. A combination of samples from 1KGP and the Human Origins [18] project were used as a reference for the projection.

## RESULTS

### Development and validation of imputation pipeline for lcWGS

Previous studies have evaluated the potential use of lcWGS in local ancestry deconvolution, complex trait association studies, and detection of rare genetic variants [4–6]. To assess the feasibility and accuracy of this approach for GPSs, we first developed an imputation pipeline that reads raw fastq sequence data and generates a vcf with imputed site information at 21.7 million sites (imputation SNP loci) (Figure 1A, B). Briefly, reads are aligned to the reference genome and filtered for duplicates and low quality. Using this BAM file, we then calculate genotype likelihoods and impute expected genotypes using 1KGP as the imputation reference panel.

To validate this imputation pipeline, we performed hcWGS and downsampling on seven samples from different 1KGP populations and a trio of Ashkenazi Jewish GIAB samples (pipeline validation data set) to varying depths of coverage from 2.0X to 0.1X (See Supplementary Table 1, Additional File 2). We used the published genotype calls for each of these samples as truth data and found that imputation accuracy was above 0.90 r^2^ for all samples at 0.5X and higher (Figure 2). As expected, this was correlated with sequencing depth, with diminishing gains observed at coverages above 1.0X. While imputation accuracy was similar across diverse populations, it was slightly reduced in the Colombian sample (HG01485), likely due to complex local ancestry related to admixture, and in the Yoruban sample (NA19240), likely due to the shorter blocks of linkage disequilibrium and higher genetic diversity in Africa [19]. Taken together, these data suggest that at sequencing depth at or above 0.5X, our pipeline has similar imputation accuracy to genotyping array-based imputation across individuals from multiple populations. As such, we set 0.5X as a quality control for success and removed samples with coverage below this threshold in subsequent analyses.

**Figure 2.**
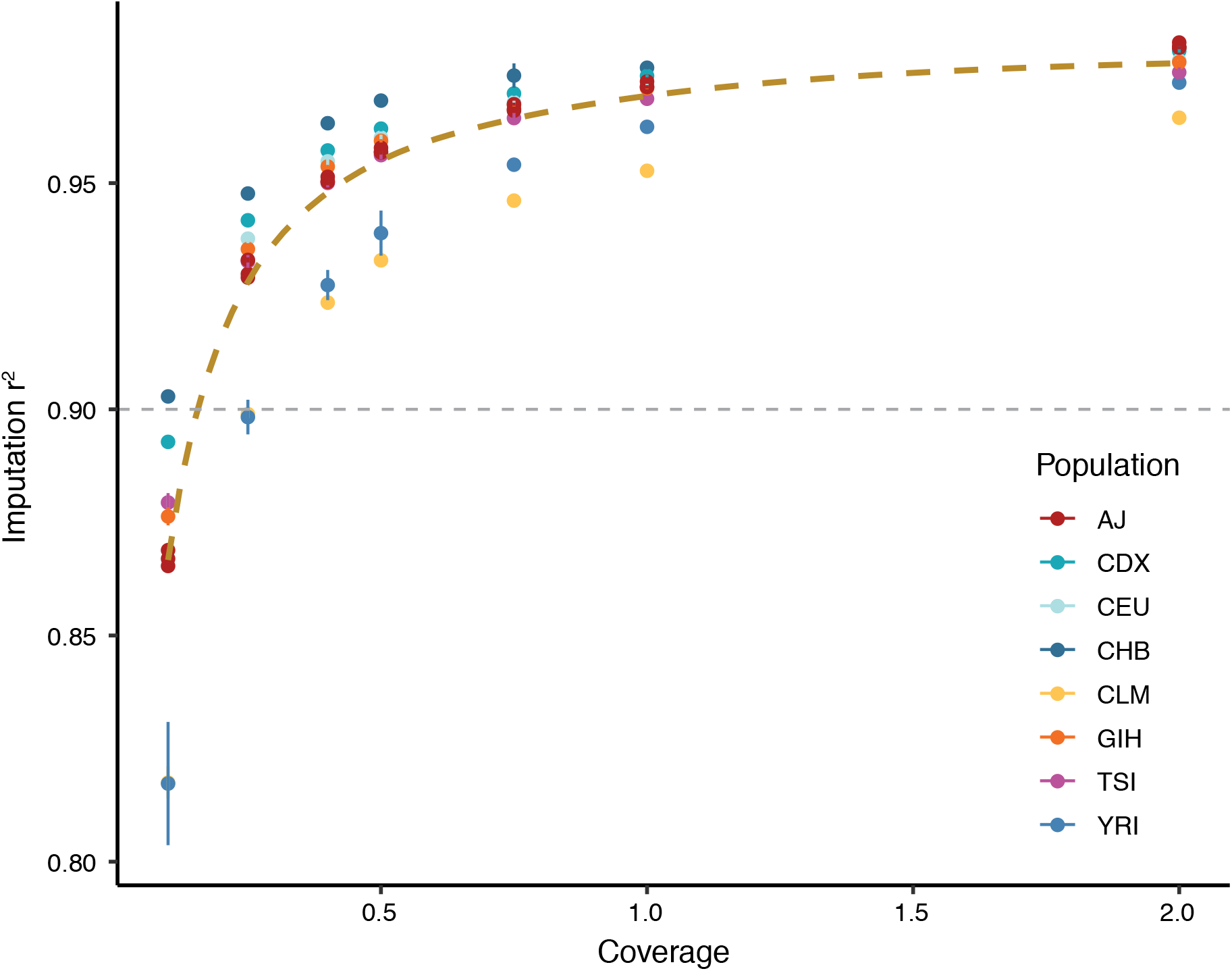
Assessment of imputation performance in the pipeline validation data set. Downsampling from 30X to 0.1X showed that lcWGS accuracy was above 0.90 r^2^ for all samples at 0.5X (n = 4 independent random seeds for each sample and coverage value; error bars are 95% confidence intervals). The thick brown dashed line is a smoothed trendline of the average imputation quality while the thin grey dashed line demonstrates previously reported imputation quality from a genotyping array (r^2^ = 0.90) [4]. AJ, Ashkenazi Jewish. CDX, Chinese Dai in Xishuangbanna, China. CEU, Utah Residents with Northern and Western European Ancestry. CHB, Han Chinese in Beijing, China. CLM, Colombians from Medellin, Colombia. GIH, Gujarati Indian from Houston, Texas. TSI, Toscani in Italia. YRI, Yoruba in Ibadan, Nigeria.

### Technical concordance between GPSs calculated from lcWGS and genotyping array

To assess the technical concordance of using lcWGS to calculate GPSs, we performed low coverage sequencing and used genotyping arrays on DNA from 184 individuals (technical concordance cohort) (Figure 1B). This concordance assessment was restricted to individuals of European ancestry to most closely align with the populations used for GPS training and validation.

We first compared the lcWGS genotype dosages with a subset of variants directly genotyped (n = 470,362) on the genotyping array to assess imputation performance. Assuming the typed loci called on the genotyping array as ‘true’, we observed an average imputation r^2^ > 0.90 at 0.5X depth for variants with global minor allele frequency (MAF) greater than 5% (see Supplementary Figure 2, Additional File 3). As expected, imputation accuracy was highest for variants with higher MAF. For lower frequency variants, we saw a reduction in imputation accuracy, as expected, with r^2^ > 0.85 for variants at 1% to 5% MAF and r^2^ > 0.80 for variants less than 1% global MAF. Taken together, this demonstrates that lcWGS has high accuracy in this test setting.

We then calculated previously published GPSs for CAD [3], BC [8], and AF [3] on each sample using genotyping array data or lcWGS data. We found that GPS_CAD_, GPS_BC_, and GPS_AF_ were highly correlated (Figure 3A-C), with the score mean (Student t-test p = 0.17) and variance (F test p = 0.91) equivalent between lcWGS and the genotyping array. The correlations of GPS_CAD_ and GPS_AF_ (r^2^ = 0.98 and r^2^ = 0.97, respectively) were slightly higher than that of GPS_BC_ (r^2^ = 0.93), which could be due to 1) the smaller number of loci in GPS_BC_ (6.6 million compared to 3820 SNPs), 2) differences in allele frequencies between SNPs with high weights, and/or 3) the fact that GPS_BC_ was trained and validated on a different genotyping array, the OncoArray, than the Axiom UK Biobank Array used in this study [8].

**Figure 3.**
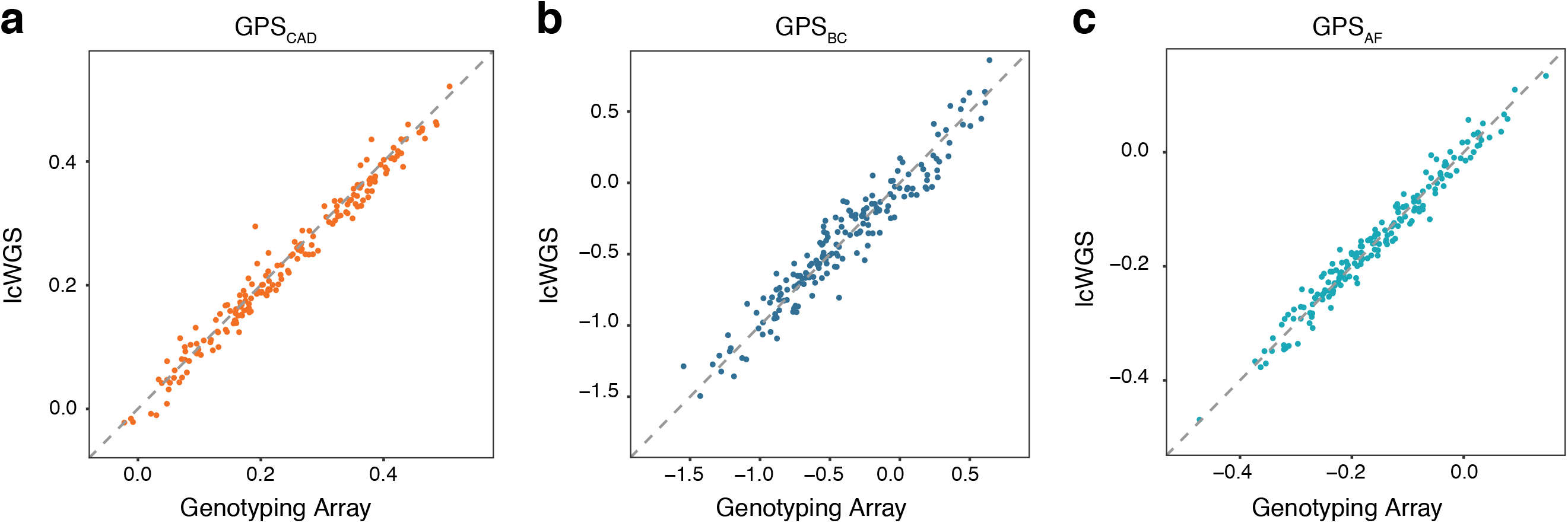
Correlation of GPSs between genotyping array and lcWGS in the technical concordance cohort. (A) GPS_CAD_ calculated using lcWGS was highly correlated (r^2^ = 0.98) with those calculated using genotyping array (n = 182). (B) GPS_BC_ calculated using lcWGS was highly correlated (r^2^ = 0.93) with those calculated using genotyping array (n = 182). (C) GPS_AF_ was highly correlated (r^2^ = 0.97) with those calculated using genotyping arrays (n = 182). x-axis is the raw GPS calculated from the genotyping array, and y-axis is the raw GPS calculated from the lcWGS data; raw GPS values are unitless. lcWGS, low coverage whole genome sequencing. GPS, genome-wide polygenic score. CAD, coronary artery disease. BC, breast cancer. AF, atrial fibrillation.

The technical concordance cohort ranged in coverage from 0.54X to 1.76X with a mean coverage of 1.24X, and we have shown that depth can impact imputation performance -- depth increases above 0.5X have a smaller but measurable effect on imputation performance (Figure 2; see Supplementary Figure 2, Additional File 3). To determine the low coverage sequencing depth required for GPS accuracy, we used SAMtools to downsample the lcWGS data in this cohort to 1.0X, 0.75X, 0.5X, 0.4X, 0.25X, and 0.1X. We found that GPS_CAD_, GPS_BC_, and GPS_AF_ are robust to lcWGS sequencing depth 0.5X and that coverages do not systematically bias GPS calculations in a specific direction (see Supplementary Figure 3 and Supplementary Figure 4, Additional File 3), indicating that samples above 0.5X with small changes in coverage variation can be combined for downstream analysis. In addition, the correlation increases logarithmically as coverage increases (see Supplementary Figure 5, Additional File 3). These data demonstrate high correlation between GPSs from lcWGS data and genotyping array in a randomly selected sample. Interestingly, correlation at 0.1X was still high enough that GPSs at this coverage may have research utility, suggesting that significant amounts of data regarding common genetic variation could be recovered from off-target reads in exome and multi-gene panel sequencing studies to allow for GPS calculation. Taken together, these data demonstrate that lcWGS provides equivalent accuracy for calculation of GPSs, with sequencing coverage as low as 0.5X.

### Assessment of imputation performance and technical concordance across diverse populations

To further assess the performance of our imputation pipeline across diverse populations, we performed lcWGS on 120 additional samples from six 1KGP populations (CHB, GIH, YRI, ASW, MXL, and PUR; see Supplementary Table 3, Additional File 1) that represent the range of ancestry observed in admixed populations (diverse ancestry data set). We compared genotypes imputed using our lcWGS pipeline to known 1KGP WGS data and found that imputation accuracy was above 0.90 r^2^ for all samples (range 0.94 - 0.97) (Figure 4A). In addition, we found that GPS calculated from lcWGS data and GPS calculated from the Phase 3 1KGP WGS data release have a high correlation, with an r^2^ value of 0.98, 0.91, and 0.98 for CAD, BC, and AF, respectively (Figure 4B-D). These results suggest that lcWGS can enable accurate imputation and calculation of GPSs in diverse populations.

**Figure 4.**
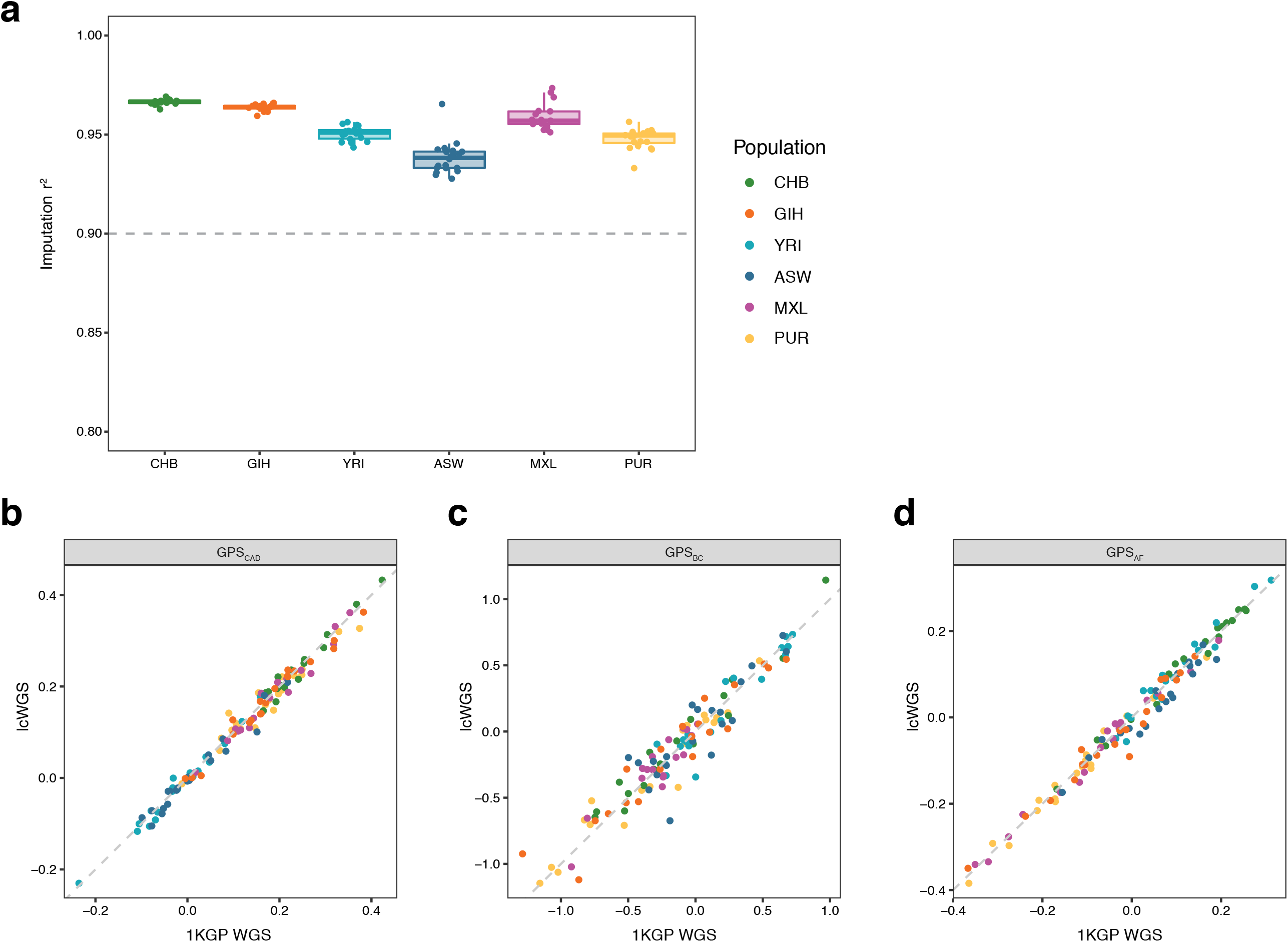
Assessment of imputation performance and technical concordance across diverse populations. (A) GPS_CAD_ calculated using lcWGS data was highly correlated with those calculated using known 1KGP data (n = 116), with all samples having a correlation coefficient above 0.90. The thin grey dashed line demonstrates previously reported imputation quality from a genotyping array (r^2^ = 0.90) [4]. (B) GPS_CAD_ calculated using lcWGS data was highly correlated (r^2^ = 0.98) with those calculated using known 1KGP data (n = 116). (C) GPS_BC_ calculated using lcWGS data was highly correlated (r^2^ = 0.91) with those calculated using known 1KGP data (n = 116). (D) GPS_AF_ was highly correlated (r^2^ = 0.98) with those calculated using known 1KGP data (n = 116). 1KGP, 1000 Genomes Project. lcWGS, low coverage whole genome sequencing. GPS, genome-wide polygenic score. CAD, coronary artery disease. BC, breast cancer. AF, atrial fibrillation.

### Association of lcWGS-calculated GPSs with disease phenotypes in a clinical cohort

Previous studies have demonstrated the association of GPSs with prevalent disease using genotyping arrays [3,8,20–22] and hcWGS [16]. To observe the performance of lcWGS-calculated GPSs in a large population, we performed low coverage sequencing on 11,502 European individuals (clinical cohort) (See Supplementary Table 2, Additional File 1) and calculated GPS_CAD_, GPS_BC_, and GPS_AF_ for each individual. Raw GPSs were normalized by taking the standardized residual of the predicted score after correction for the first 10 PCAs (see Supplementary Figure 6, Additional File 3) [16,23]. First, we note that there are no major outliers (defined as a z-score greater than 5) in GPS_CAD_, GPS_BC_, and GPS_AF_ and that the normalized scores formed an approximately normal distribution for each (see Supplementary Figure 7, Additional File 3). Each of the GPSs were strongly associated with self-reported history of disease, with effect estimates comparable to prior reports using genotyping arrays to calculate GPS -- GPS_CAD_ (OR per standard deviation = 1.59 (1.32 - 1.92), n = 11,010), GPS_BC_ (OR per standard deviation = 1.56 (1.45 - 1.68); n = 8722), and GPS_AF_ (OR per standard deviation = 1.28 (1.12 - 1.46); n = 10,303) (Figure 5).

**Figure 5.**
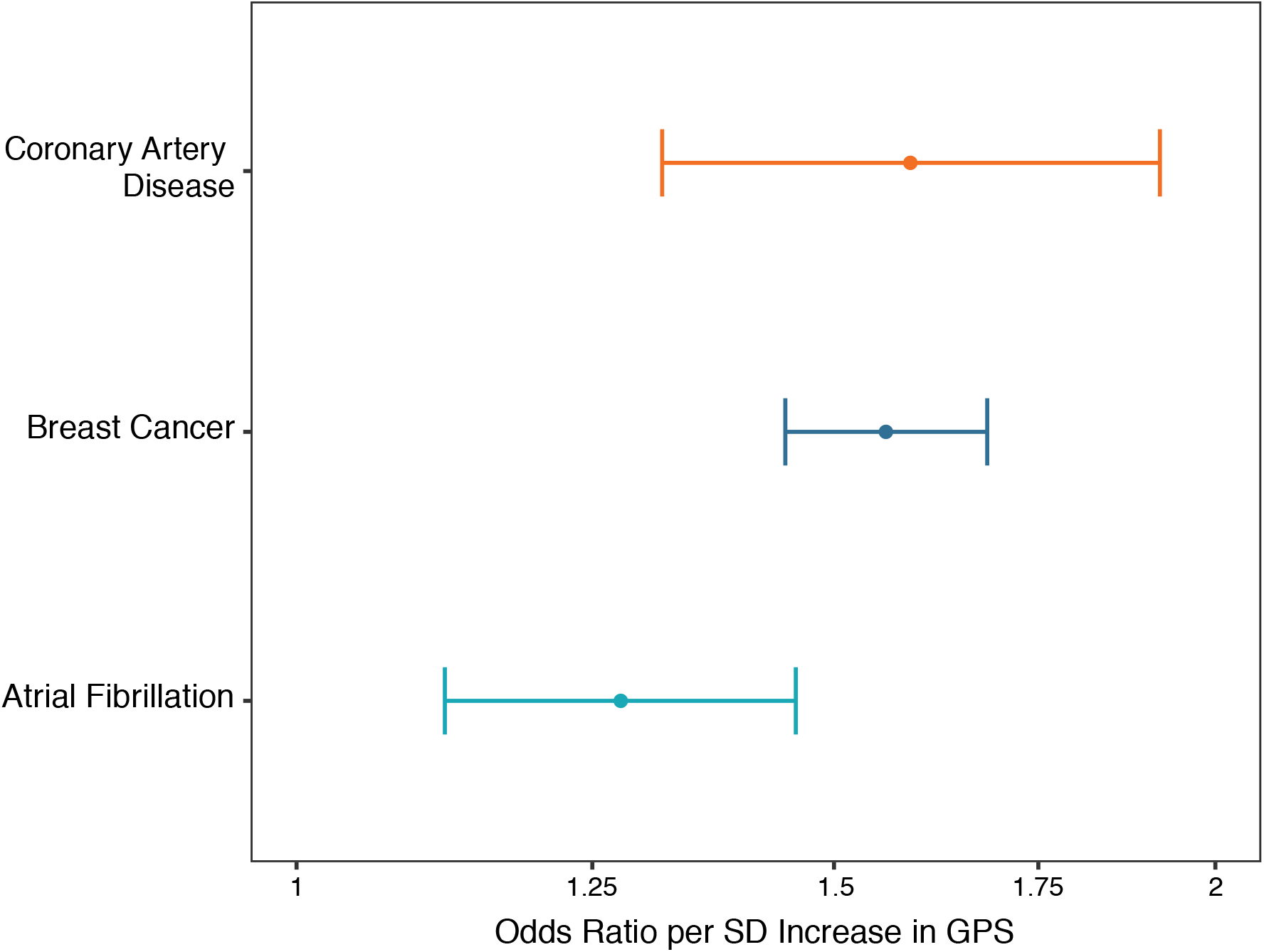
Association of lcWGS-calculated GPSs with disease phenotypes in the clinical cohort. lcWGS-calculated GPS_CAD_ was associated with personal history of CAD (OR = 1.589 (1.32 - 1.92), n = 11,010, p = 1.32 × 10^−6^). GPS_CAD_ was adjusted for age and sex. lcWGS-calculated GPS_BC_ was associated with personal history of BC (OR = 1.56 (1.45 - 1.68); n = 8,722, p = 1.0 × 10^−16^). GPS_BC_ was calculated only for females and adjusted for age at menarche. lcWGS-calculated GPS_AF_ was associated with personal history of AF (OR = 1.277 (1.12 - 1.46); n = 10,303, p = 0.000292). GPS_AF_ was adjusted for age and sex. lcWGS, low coverage whole genome sequencing. GPS, genome-wide polygenic score. CAD, coronary artery disease. BC, breast cancer. AF, atrial fibrillation.

Previous studies have noted significantly increased disease prevalence among individuals in the extreme tails of the GPS distribution when compared to the remainder of the population [3,8]. We replicated this observation by assessing the prevalence of disease in the highest 5% of the GPS distribution for each of the three diseases, noting odds ratios of 4.5 (2.62 - 7.74), 2.62 (2.04 - 3.36), and 1.96 (1.24 - 3.11) for GPS_CAD_, GPS_BC_, and GPS_AF_, respectively.

Area under the curve (AUC) is an additional metric used to assess the ability of a given risk factor to discriminate between affected cases and disease-free controls. When only the GPS was included in the prediction model, GPS_CAD_ had an AUC of 0.60, GPS_BC_ had an AUC of 0.63, and GPS_AF_ had an AUC of 0.57. The additional inclusion of age and sex increased the AUCs to 0.86 for GPS_CAD_, 0.78 for GPS_BC_, and 0.78 for GPS_AF_. For each of these three common, complex diseases, the magnitude of associations with clinical disease and AUC metrics were consistent with previous publications [3,8]. Taken together, these results suggest that lcWGS-calculated GPSs can accurately stratify risk with comparable accuracy to previously published GPS-disease associations calculated on the basis of genotyping array data.

## DISCUSSION

For the past two decades, genotyping array-based GWAS and imputation have been the driving force in our discovery of genetic loci predictive of disease and derivation and calculation of GPSs. In this study, we developed and validated an imputation pipeline to calculate GPSs from variably downsampled hcWGS and lcWGS data sets. While the efficiency of lcWGS has been reported for other applications of statistical genetics [4–6], we demonstrate that lcWGS achieves similar technical concordance as the Axiom UK Biobank Array by Affymetrix for determining GPSs. Furthermore, the imputation r^2^ from lcWGS was greater than 90%, which is similar to the imputation accuracy reported from other commercially-available genotyping arrays [24]. Taken together, these data suggest that lcWGS has comparable accuracy to genotyping arrays for assessment of common variants and subsequent calculation of GPSs.

Our finding that lcWGS can be used for accurate genotyping and imputation of common genetic variants has implications for the future of genomic research and medicine. Currently, disease GWAS are performed using a variety of genotyping arrays that are designed to target specific sets of genes or features, reducing imputation quality in regions that are not targeted [25]. lcWGS enables less biased imputation than genotyping arrays by not pre-specifying the genetic content that is included for assessment, as is necessary for genotyping arrays. Because initial GWAS focused on populations with high homogeneity to reduce noise and increase fit of risk stratification, many genotyping arrays were designed to capture common genetic variants based on the linkage disequilibrium structure in European populations [26]. However, this ascertainment bias reduces the imputation performance from genotyping array data in diverse populations [27–29]. Imputation from lcWGS data reduces this bias by including all SNPs observed in 1KGP populations as potential predictors. The effects of SNP selection bias are also not equivalent across genotyping arrays, and therefore variants included in a GPS trained and validated on one genotyping array may not be as predictive on another genotyping array [30]. lcWGS systematically surveys variants independent of SNP selection bias and thus provides one approach to overcome this issue. Our findings here demonstrate that GPSs trained and validated on different genotyping arrays are transferable to lcWGS-calculated GPS.

Furthermore, as new genetic associations are discovered, lcWGS can be re-analyzed with ever more inclusive sets of known SNPs, further reducing SNP selection bias and advancing the study and understanding of the genetic contributions to disease. In contrast, genotyping arrays are static and cannot be easily updated or changed without designing a *de novo* platform.

lcWGS also has the potential to easily integrate into current clinical sequencing pipelines. In contrast to genotyping arrays, which require investment in separate laboratory technology, lcWGS can be performed on the same platform as current hcWGS or targeted multi-gene panel clinical testing. The ease of combining these two pathways could help to drive GPS adoption into clinical practice and can likely be achieved at a cost comparable to genotyping arrays [4]. As the cost of next generation sequencing continues to decrease, the cost of lcWGS will also continue to decrease.

This study should be interpreted in the context of potential limitations. First, the imputation accuracy observed in our analysis may have been limited by the reference panel size. Future efforts using an even larger reference panel may lead to further improved imputation accuracy, particularly for variants with allele frequency less than 1% [24]. Second, while lcWGS may ultimately enable derivation of GPSs with improved predictive accuracy or ethnic transferability, this was not explicitly explored here. Rather, we demonstrate the feasibility and accuracy of using lcWGS of calculating GPSs published in previous studies. Third, disease phenotypes in our clinical cohort were based on individual self-report rather than review of health records. However, several studies have shown that self-reported personal history data have high concordance with data reported by a healthcare provider or electronic health records [31–34], and any inaccuracies would be expected to bias GPS-disease associations to the null.

## CONCLUSIONS

In conclusion, this work establishes lcWGS as an alternative approach to genotyping arrays for common genetic variant assessment and GPS calculation -- providing comparable accuracy at similar cost while also overcoming the ascertainment bias inherent to variant selection in genotyping array design.

## LIST OF ABBREVIATIONS

GPS: genome-wide polygenic score
lcWGS: low coverage whole genome sequencing
CAD: coronary artery disease
BC: breast cancer
AF: atrial fibrillation
1KGP: 1000 Genomes Project
GIAB: Genome in a Bottle
Indel: insertion-deletion
BWA: Burrows-Wheeler Aligner
IBS: Identity-by-State
PC: principal components
PCA: PC analysis
MAF: minor allele frequency
AUC: area under the curve

## DECLARATIONS

### Ethics approval and consent to participate

All individuals in the technical concordance cohort and clinical cohort gave electronic informed consent to have their de-identified information and sample used in anonymized studies (Western Institutional Review Board, #20150716).

### Consent for publication

All individuals in the technical concordance cohort and clinical cohort gave electronic informed consent that Color may author publications using non-aggregated, de-identified information, either on its own or in collaboration with academic or commercial third parties.

### Availability of data and material

The technical concordance and clinical cohort data are not publicly available given the potential to compromise research participant privacy or consent.

1KGP, http://www.internationalgenome.org/

GIAB, ftp://ftp-trace.ncbi.nlm.nih.gov/giab/ftp/release/AshkenazimTrio/

Samtools/Bcftools, http://www.htslib.org/

BEAGLE, https://faculty.washington.edu/browning/beagle/beagle.html

FastNGSAdmix, http://www.popgen.dk/software/index.php/FastNGSadmix

### Authors’ contributions

JRH, GM, AYZ, and AVK designed the overall study. JRH, CLN, GM, AYZ, SK, and AVK contributed to data acquisition and analysis. JRH, CLN, GM, AYZ, SK, and AVK drafted or critically revised the manuscript for important intellectual content. AYZ and AVK are the guarantors of this work and, as such, have full access to all of the data in the study and take responsibility for the integrity of the data and the accuracy of the data analysis.

## Acknowledgements

We would like to thank Will Stedden, Carmen Lai, and Anjali D. Zimmer for helpful discussions and Justin Lock, Alok Sabnis, and Valerie Ngo for laboratory support and sample processing.

## ADDITIONAL FILES

Additional File 1

Homburger et al Additional File 1, PDF

Supplementary tables and legends

Additional File 2

Homburger et al Additional File 2, PDF

Supplementary Methods

Additional File 3

Homburger et al Additional File 3, PDF

Supplementary figures and legends

