## Additional Files for "Low coverage whole genome sequencing enables accurate assessment of common variants and calculation of genome-wide polygenic scores"

**ADDITIONAL FILE 1**

**Supplementary tables and legends**

Supplementary Table 1. Samples in the pipeline validation data set.

| Sample | Population code | Population description | Super population code |
|---|---|---|---|
| NA24385 | AJ | Ashkenazi Jewish Trio, Son | AJ |
| NA24143 | AJ | Ashkenazi Jewish Trio, Mother | AJ |
| NA24149 | AJ | Ashkenazi Jewish Trio, Father | AJ |
| HG02155 | CDX | Chinese Dai in Xishuangbanna, China | EAS |
| NA12878 | CEU | Utah Residents with Northern and Western European Ancestry | EUR |
| HG00663 | CHB | Southern Han Chinese | EAS |
| HG01485 | CLM | Colombians from Medellin, Colombia | AMR |
| NA21144 | GIH | Gujarati Indian from Houston, Texas | SAS |
| NA20510 | TSI | Toscani in Italia | EUR |
| NA19420 | YRI | Yoruba in Ibadan, Nigeria | AFR |

AJ, Ashkenazi Jewish. EAS, East Asian. EUR, European. AMR, Ad Mixed American. SAS, South Asian. AFR, African.

Supplementary Table 2. Demographics of technical concordance cohort and clinical cohort.

| | | Technical concordance cohort | | Clinical cohort | |
|---|---|---|---|---|---|
| | | Individuals (n) | Population | Individuals (n) | Population |
| Total | | 182* | 100% | 11,502 | 100% |
| Gender | Female | 127 | 69.8% | 9529 | 82.8% |
| | Male | 55 | 30.2% | 1973 | 17.1% |
| Age (Years) | 18-30 | 6 | 3.3% | 1041 | 9.1% |
| | 31-40 | 16 | 8.8% | 2410 | 21.0% |
| | 41-50 | 37 | 20.3% | 2633 | 22.9% |
| | 51-65 | 65 | 35.7% | 3907 | 34.0% |
| | 65+ | 58 | 31.9% | 1511 | 13.1% |
| Personal History | CAD | 61 | 33.5% | 126 | 1.1% |
| | BC | 18† | 14.1% | 828† | 8.6% |
| | AF | 0 | 0.0% | 239 | 2.1% |

*Excludes two individuals who failed genotyping and low coverage whole genome sequencing.

†Females only. CAD, coronary artery disease. BC, breast cancer. AF, atrial fibrillation.

Supplementary Table 3. Samples in the diverse ancestry data set.

| Population code | Population description | Super population code | Samples | | | |
|---|---|---|---|---|---|---|
| CHB | Han Chinese in Beijing, China | EAS | NA18642<br>NA18757<br>NA18564<br>NA18609<br>NA18597 | NA18619<br>NA18563<br>NA18749<br>NA18621<br>NA18534 | NA18544<br>NA18555<br>NA18645<br>NA18560<br>NA18577 | NA18631<br>NA18634<br>NA18543<br>NA18582<br>NA18626 |
| YRI | Yoruba in Ibadan, Nigeria | AFR | NA18501<br>NA18502<br>NA18505<br>NA18507<br>NA18508 | NA18516<br>NA18519<br>NA18861<br>NA18867<br>NA18868 | NA18873<br>NA18910<br>NA18917<br>NA19095<br>NA19114 | NA19117<br>NA19129<br>NA19137<br>NA19143<br>NA19147 |
| GIH | Gujarati Indian from Houston, Texas | SAS | NA20870<br>NA20910<br>NA20886<br>NA20900<br>NA20845 | NA20903<br>NA21142<br>NA21141<br>NA21105<br>NA21088 | NA20899<br>NA21133<br>NA21137<br>NA21098<br>NA21114 | NA21115<br>NA21103<br>NA21125<br>NA21126<br>NA21094 |
| ASW | Americans of African Ancestry in Southwest USA | AFR | NA20314<br>NA19921<br>NA19625<br>NA19914<br>NA19818 | NA19834<br>NA20359<br>NA19700<br>NA19701<br>NA20348 | NA20356<br>NA19908<br>NA19916<br>NA20127<br>NA20291 | NA19707<br>NA20342<br>NA20317<br>NA20351<br>NA19819 |
| MXL | Mexican Ancestry from Los Angeles, USA | AMR | NA19729<br>NA19728<br>NA19731<br>NA19732<br>NA19678 | NA19679<br>NA19651<br>NA19676<br>NA19794<br>NA19725 | NA19770<br>NA19776<br>NA19661<br>NA19723<br>NA19771 | NA19761<br>NA19786<br>NA19762<br>NA19785<br>NA19759 |
| PUR | Puerto Ricans from Puerto Rico | AMR | HG01241<br>HG01188<br>HG01051<br>HG01167<br>HG01049 | HG01171<br>HG01101<br>HG00734<br>HG01182<br>HG01183 | HG00740<br>HG01048<br>HG01204<br>HG01098<br>HG01248 | HG00551<br>HG01067<br>HG01104<br>HG01197<br>HG00640 |

EAS, East Asian. AFR, African. SAS, South Asian. AMR, Ad Mixed American.

**ADDITIONAL FILE 2**

**Supplementary Methods**

The multi-gene NGS panel test analyzed genes that have been associated with an elevated risk of hereditary cancer or hereditary heart conditions. These genes were selected based on published evidence of association and technical feasibility using the methods described. Analysis, variant calling, and reporting focused on the complete coding sequence and adjacent intronic sequence of the primary transcript(s), unless otherwise indicated*.

For hereditary cancer, these genes are *APC, ATM, BAP1, BARD1, BMPR1A, BRCA1, BRCA2, BRIP1, CDH1, CDK4*, CDKN2A* (p14ARF and p16INK4a), *CHEK2, EPCAM*, GREM1*, MITF*, MLH1, MSH2, MSH6, MUTYH, NBN, PALB2, PMS2*, POLD1*, POLE*, PTEN, RAD51C, RAD51D, SMAD4, STK11,* and *TP53*. Exons 12-15 of *PMS2* cannot be reliably assessed with standard target enrichment protocols. For the *CDK4*, *MITF*, *POLD1,* and *POLE*, the elevated risk of cancer is associated with distinct functional genomic regions; therefore, the complete coding sequences of these genes were not reported, but instead only the following regions: *CDK4* - chr12:g.58145429-58145431 (codon 24), *MITF* - chr3:g.70014091 (including c.952G>A), *POLD1* - chr19:g.50909713 (including c.1433G>A) and *POLE* - chr12:g.133250250 (including c.1270C>G). For *EPCAM*, only deletions including the 3' end of the gene (exons 8 and/or 9) were reported. *GREM1* was only analyzed for duplications in the upstream regulatory region.

For hereditary heart conditions, these genes are *ACTA2*, *ACTC1*, *APOB**, *COL3A1*, *DSC2*, *DSG2*, *DSP*, *FBN1*, *GLA*, *KCNH2**, *KCNQ1**, *LDLR**, *LMNA*, *MYBPC3*, *MYH7**, *MYH11*, *MYL2*, *MYL3*, *PCSK9*, *PKP2*, *PRKAG2*, *RYR2*, *SCN5A*, *SMAD3*, *TGFBR1**, *TGFBR2*, *TMEM43*,

*TNNI3*, *TNNT2*, and *TPM1*. *APOB* exon 1, *KCNH2* exon 4, *KCNQ1* exon 1 and *TGFBR1* exon 1 cannot be reliability assessed with standard target enrichment protocols. For the *LDLR* promoter region, the detection of deletions, duplications, and complex structural rearrangements may be limited. For the *LDLR* promoter region, the detection of deletions, duplications, and complex structural rearrangements may be limited.

Variants were classified according to the American College of Medical Genetics and Genomics 2015 guidelines for sequence variant interpretation, and all variant classifications were signed out by a board certified medical geneticist or pathologist.

**ADDITIONAL FILE 3**

**Supplementary figures and legends**

Supplementary Figure 1. Association of lcWGS time and accuracy for samples in the pipeline validation data set at 1.0X coverage. (A) Imputation time increased as the size of the reference panel increased. (B) Imputation accuracy increased as the size of the reference panel increased, with less improvement after a panel size of 250. lcWGS, low coverage whole genome sequencing.

Supplementary Figure 2. Imputation performance of the pipeline compared to genotyping array for different allele frequencies. Imputation quality was highest for variants above 5% global MAF but was reduced for variants with lower allele frequencies. MAF, minor allele frequency.

Supplementary Figure 3. Correlation of GPSs between genotyping array and lcWGS at different coverage depths in the technical concordance cohort. (A, B) Downsampling from 1.0X to 0.1X showed that $GPS_{CAD}$ calculated using lcWGS was highly correlated with the genotyping array at 1.0X to 0.5X coverage but decreased at 0.1X (n = 182, two independent random seeds). (C, D) Downsampling from 1.0X to 0.1X showed that $GPS_{BC}$ calculated using lcWGS was highly correlated with the genotyping array at 1.0X to 0.5X coverage but decreased at 0.1X (n = 182, two independent random seeds). (D, E) Downsampling from 1.0X to 0.1X showed that $GPS_{AF}$ calculated using lcWGS was highly correlated with the genotyping array at 1.0X to 0.5X coverage but decreased at 0.1X (n = 182, two independent random seeds). x-axis is the raw GPS calculated from the genotyping array, and y-axis is the raw GPS calculated from the lcWGS data; raw GPS values are unitless. lcWGS, low coverage whole genome sequencing.

GPS, genome-wide polygenic score. CAD, coronary artery disease. BC, breast cancer. AF, atrial fibrillation.

Supplementary Figure 4. Correlation of $GPS_{CAD}$ between genotyping array and lcWGS at different coverage depths in the technical concordance cohort when removing individuals who were suspected to have a high $GPS_{CAD}$. (A) The analyzed technical concordance cohort included 59 individuals who were suspected to have a high $GPS_{CAD}$ based on targeted panel and off target sequencing data, which could artificially inflate the observed correlation above what would be seen in a randomly selected sample. $GPS_{CAD}$ remained highly correlated after removing those individuals who were suspected to have a high $GPS_{CAD}$ based on targeted panel and off target sequencing data ($r^2$ = 0.97, n = 123). (B, C) Downsampling from 1.0X to 0.1X showed that $GPS_{CAD}$ calculated using lcWGS was highly correlated with the genotyping array at 1.0X to 0.5X coverage but decreased at 0.1X (n = 123, two independent random seeds). x-axis is the raw GPS calculated from the genotyping array, and y-axis is the raw GPS calculated from the lcWGS data; raw GPS values are unitless. lcWGS, low coverage whole genome sequencing. GPS, genome-wide polygenic score. CAD, coronary artery disease.

Supplementary Figure 5. Concordance of GPS calculated at different coverages using different sampling seeds in the technical concordance cohort. GPS concordance is > 0.80 $r^2$ when samples were sequenced at or above 0.5X coverage (n = 182) ($GPS_{CAD}$ p = 0.004, $GPS_{BC}$ p = 0.0001, $GPS_{AF}$ p = 0.004). GPS, genome-wide polygenic score. CAD, coronary artery disease. BC, breast cancer. AF, atrial fibrillation.

Supplementary Figure 6. First two principal components of ancestry. Raw GPSs were normalized by taking the standardized residual of the predicted score after correction for the first 10 principal components (PC) of ancestry. Grey points correspond to reference samples from the 1KGP and Human Origins. Black points represent individuals in the clinical cohort in this study. GPS, genome-wide polygenic score. 1KGP, 1000 Genome Project.

Supplementary Figure 7. Distribution of GPSs in the clinical cohort. (A) The distribution of $GPS_{CAD}$ was approximately normal in 11,010 individuals ($p = 1.32 \times 10^{-6}$). (B) The distribution of $GPS_{BC}$ was approximately normal in 8,722 individuals ( $p = 1.0 \times 10^{-16}$). (C) The distribution of $GPS_{AF}$ was approximately normal in 10,303 individuals ($p = 0.000292$). GPS, genome-wide polygenic score. CAD, coronary artery disease. BC, breast cancer. AF, atrial fibrillation.

### Figure S1

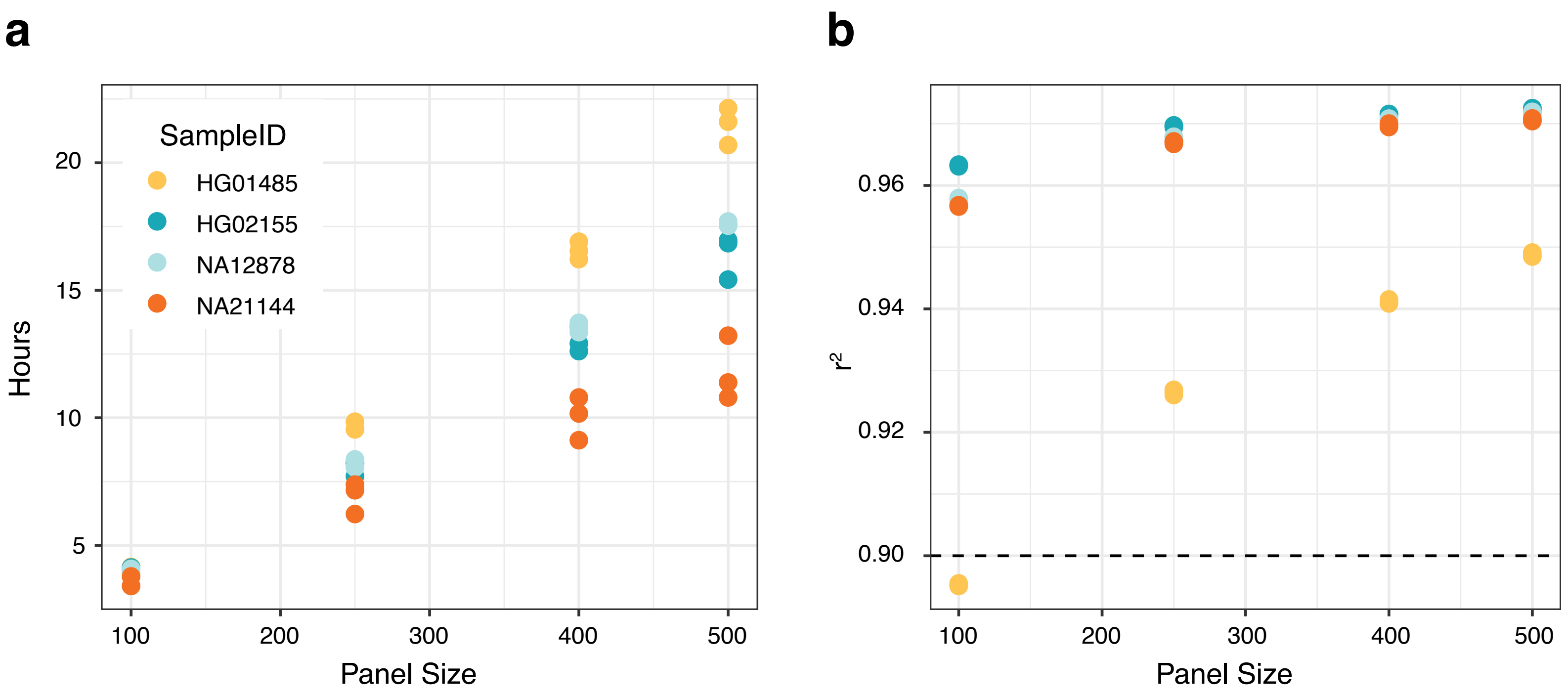
a
SampleID
HG01485
HG02155
NA12878
NA21144
Hours
20
15
10
5
100
200
300
400
500
Panel Size
b
$r^2$
0.96
0.94
0.92
0.90
100
200
300
400
500
Panel Size

### Figure S2

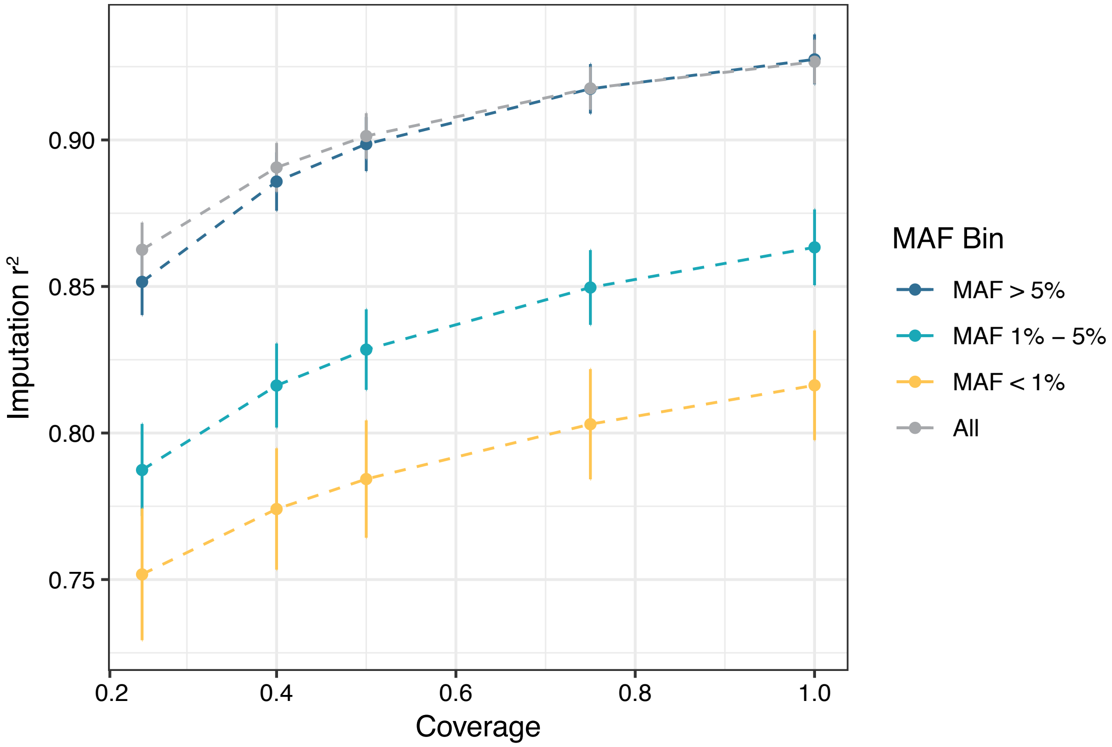
Imputation r²
0.90
0.85
0.80
0.75
0.2
0.4
0.6
0.8
1.0
Coverage
MAF Bin
MAF > 5%
MAF 1% – 5%
MAF < 1%
All

### Figure S3

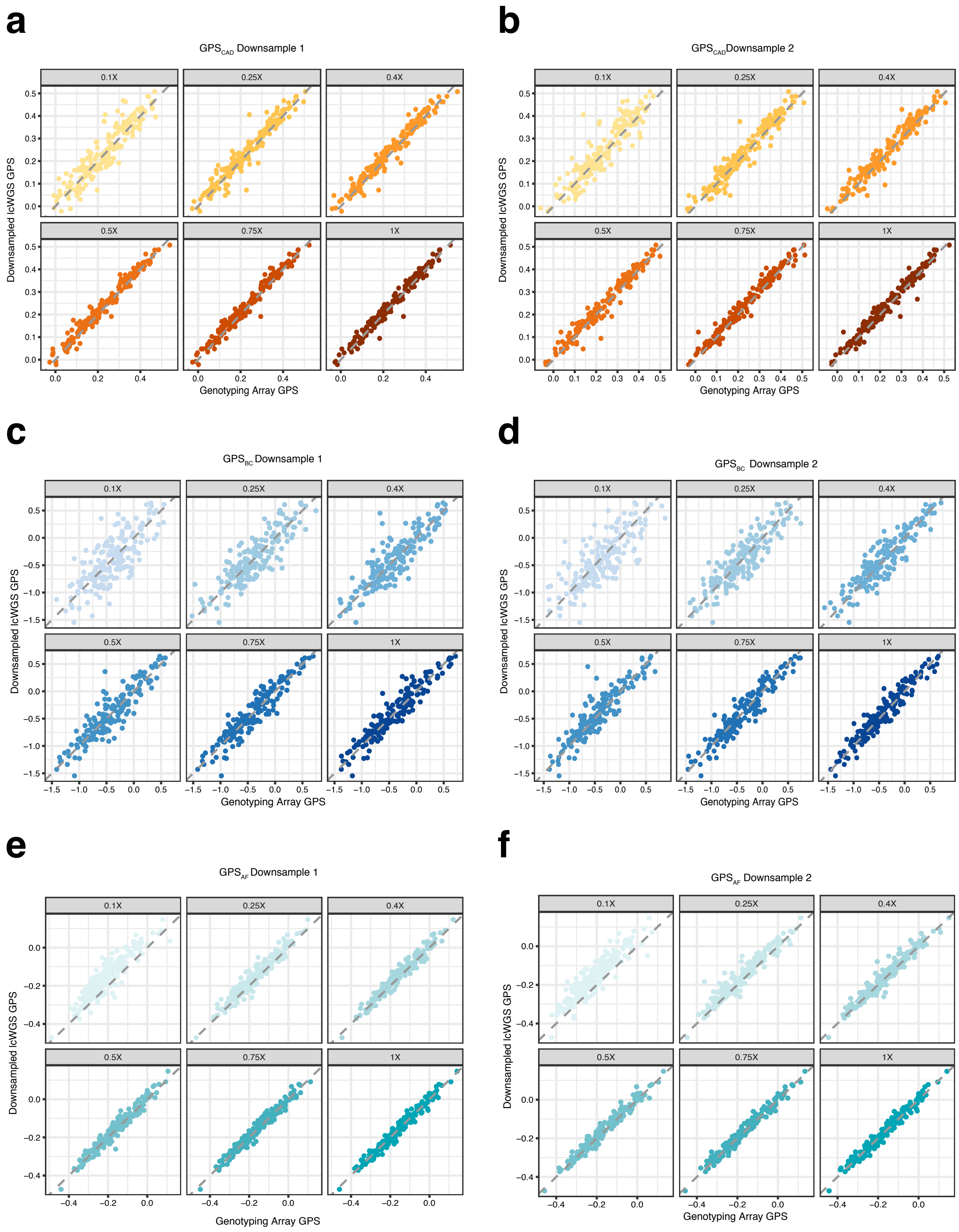
a
GPS$_{CAD}$ Downsample 1
b
GPS$_{CAD}$Downsample 2
0.1X
0.25X
0.4X
0.5X
0.75X
1X
Downsampled lcWGS GPS
Genotyping Array GPS
c
GPS$_{BC}$ Downsample 1
d
GPS$_{BC}$ Downsample 2
e
GPS$_{AF}$ Downsample 1
f
GPS$_{AF}$ Downsample 2

### Figure S4

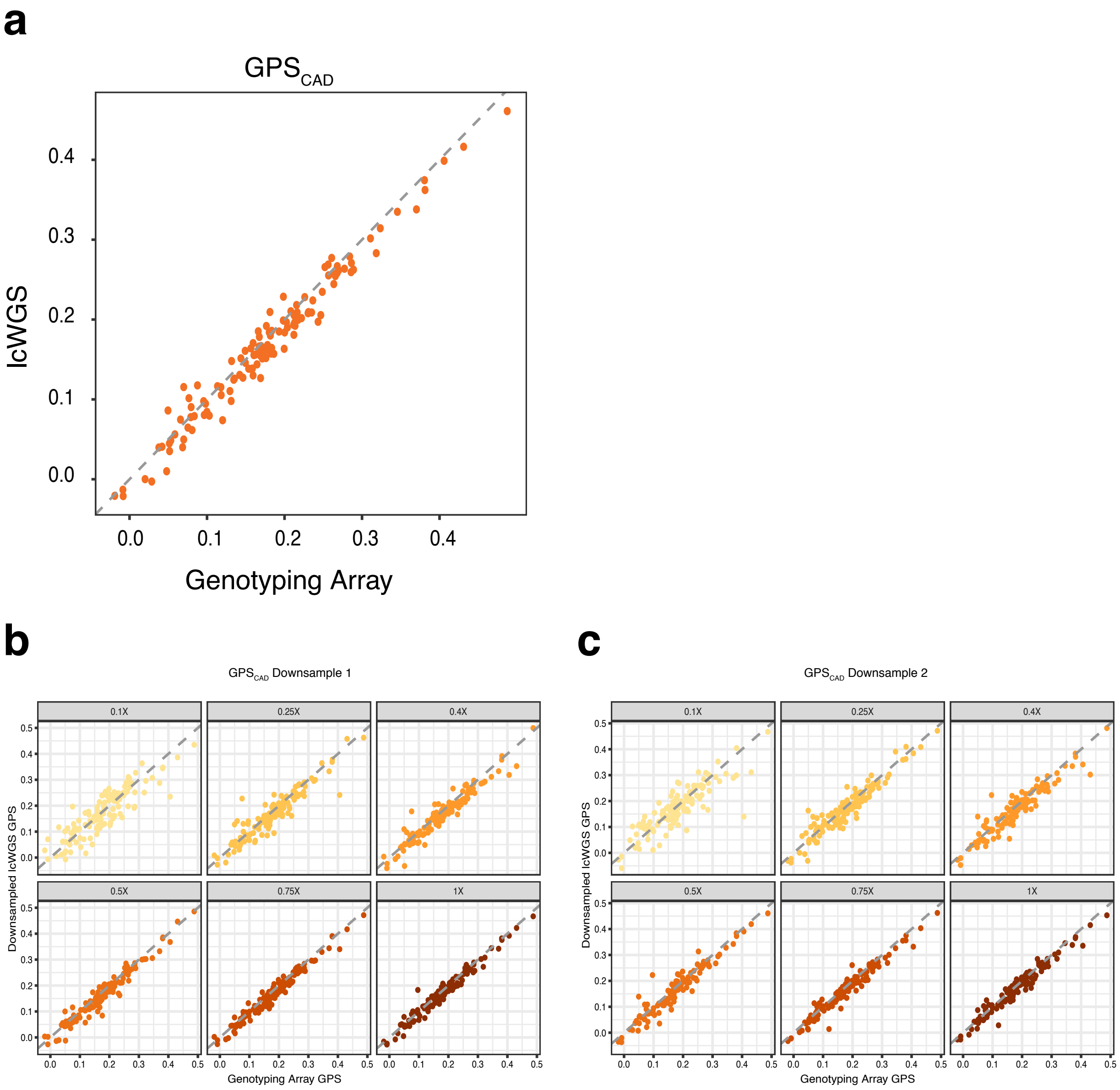
a
GPS CAD
lcWGS
Genotyping Array
0.0
0.1
0.2
0.3
0.4
b
GPS CAD Downsample 1
0.1X
0.25X
0.4X
0.5X
0.75X
1X
Downsampled lcWGS GPS
Genotyping Array GPS
c
GPS CAD Downsample 2
Downsampled lcWGS GPS
Genotyping Array GPS

### Figure S5

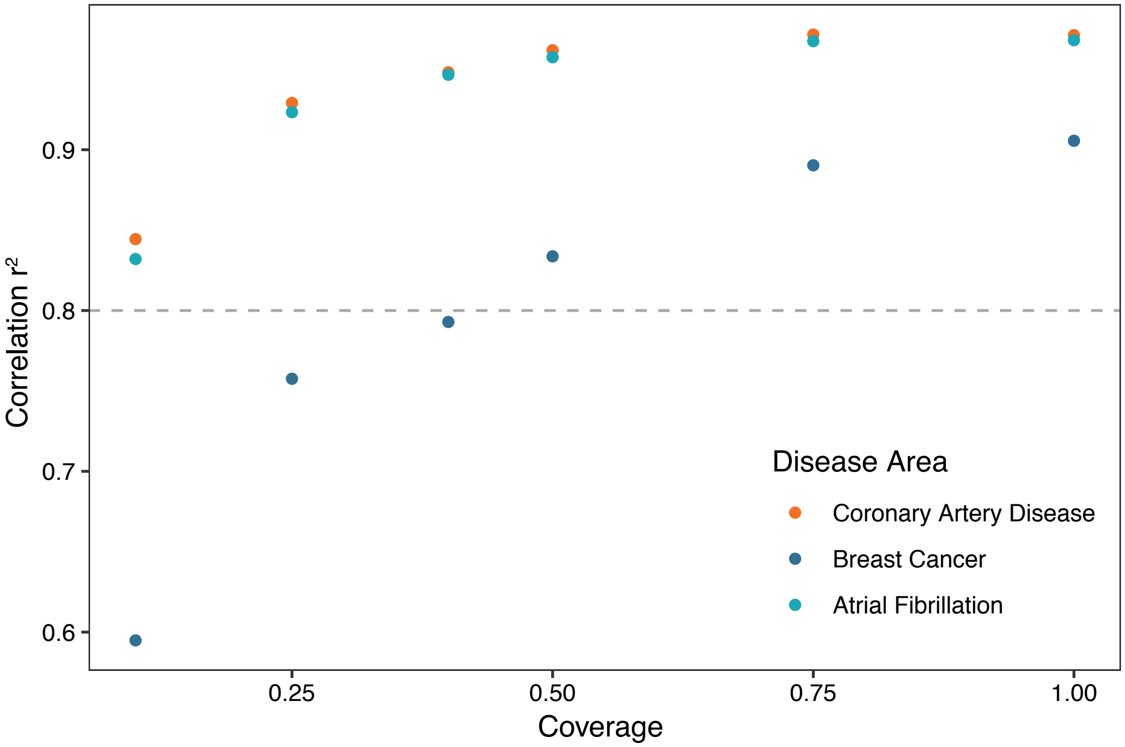

Correlation r²
0.9
0.8
0.7
0.6
0.25
0.50
0.75
1.00
Coverage
Disease Area
Coronary Artery Disease
Breast Cancer
Atrial Fibrillation

### Figure S6

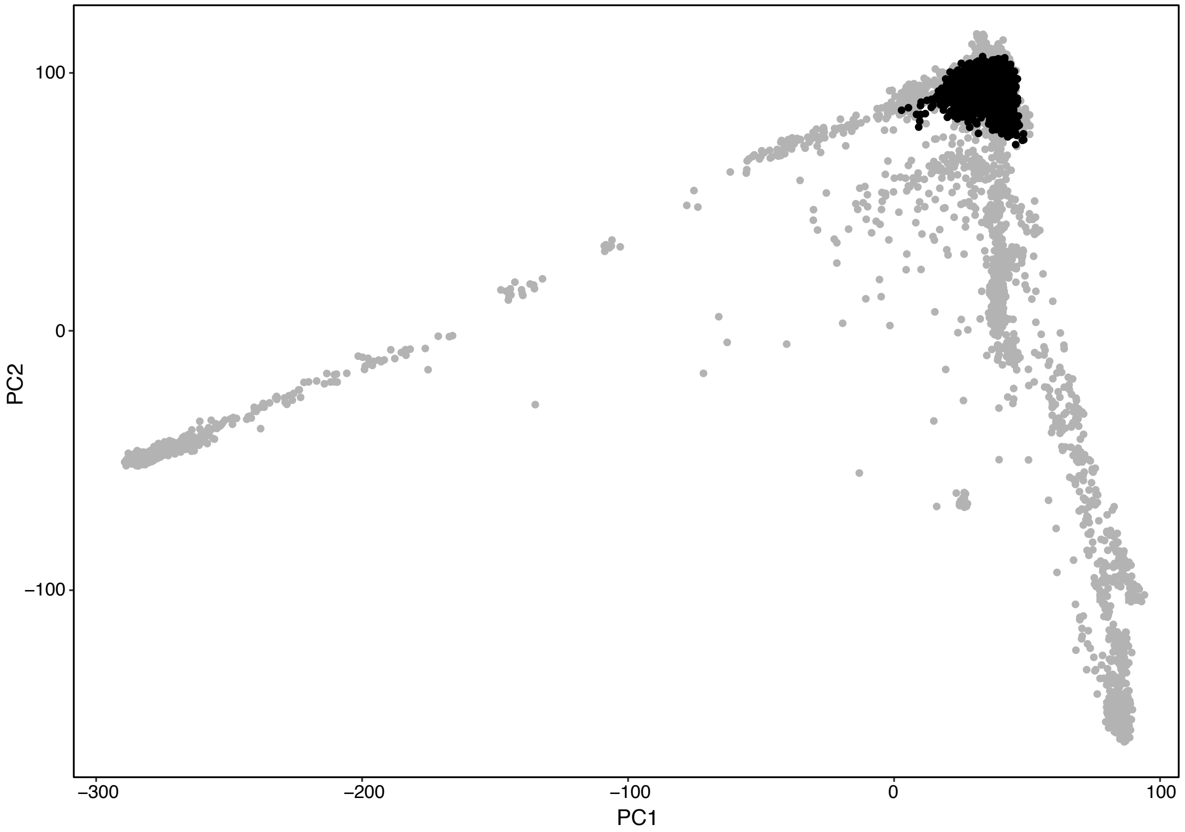

### Figure S7

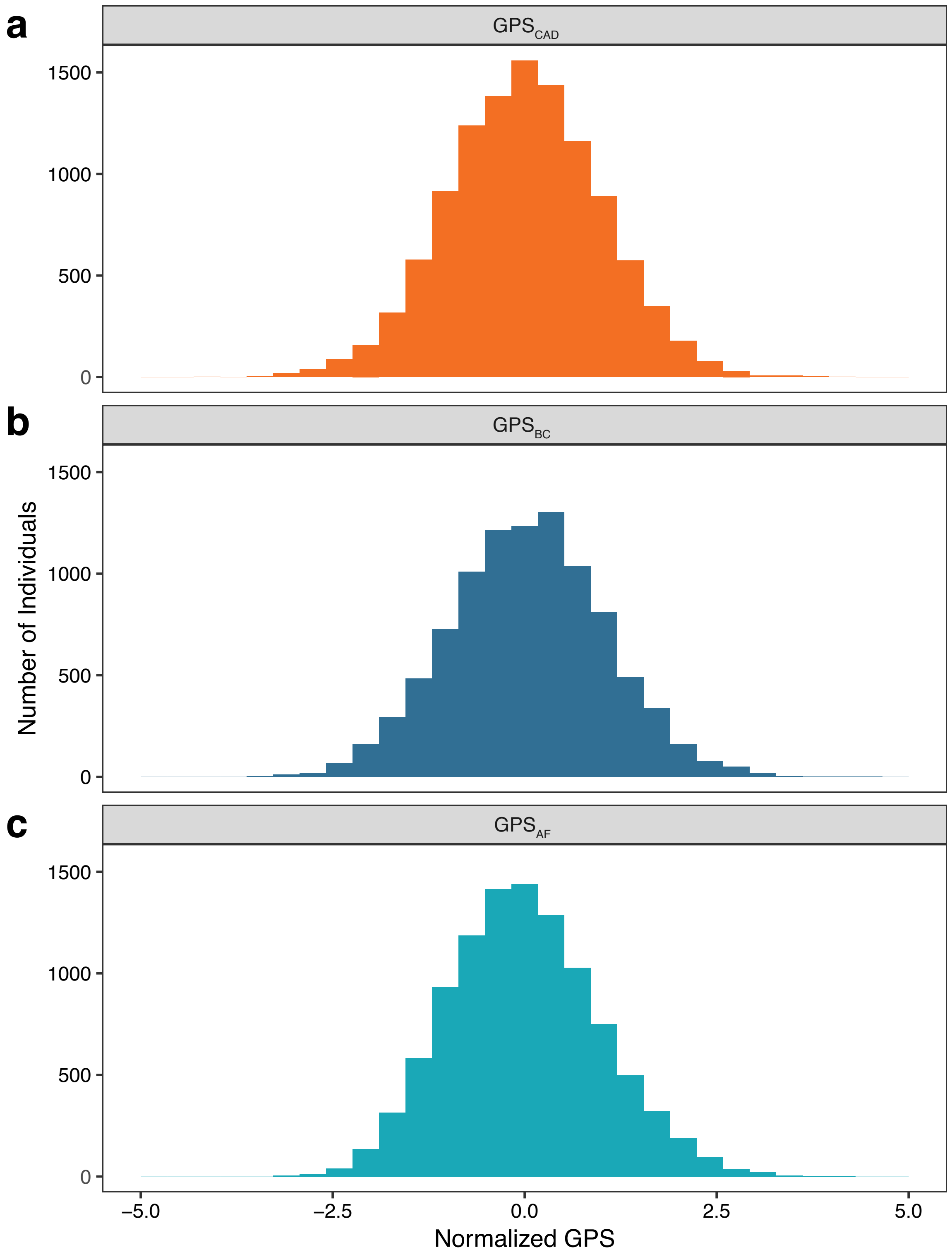
a
b
c
GPS$_{CAD}$
GPS$_{BC}$
GPS$_{AF}$
1500
1000
500
0
Number of Individuals
−5.0
−2.5
0.0
2.5
5.0
Normalized GPS
